# Phencyclidine disrupts neural coordination and cognitive control by dysregulating translation

**DOI:** 10.1101/2022.11.18.517075

**Authors:** Eun Hye Park, Hsin-Yi Kao, Hussam Jourdi, Milenna van Dijk, Simón Carrillo-Segura, Kayla W. Tunnell, Jeffrey Gutierrez, Emma J. Wallace, Matthew Troy-Regier, Basma Radwan, Edith Lesburguères, Juan Marcos Alarcon, André A Fenton

## Abstract

**Background:** Phencyclidine (PCP) causes psychosis, is abused with increasing frequency, and was extensively used in antipsychotic drug discovery. PCP discoordinates hippocampal ensemble action potential discharge and impairs cognitive control in rats, but how this uncompetitive N-methyl-D-aspartate receptor (NMDAR) antagonist impairs cognition remains unknown.

**Methods:** The effects of PCP were investigated i) on hippocampal CA1 ensemble action potential discharge *in vivo* in urethane-anesthetized rats and during awake behavior in mice; ii) on synaptic responses in *ex vivo* mouse hippocampus slices; iii) in mice on a hippocampus-dependent active place avoidance task that requires cognitive control; and iv) on activating the molecular machinery of translation in acute hippocampus slices. Mechanistic causality was assessed by comparing the PCP effects to the effects of inhibitors of protein synthesis, group-1-type metabotropic glutamate receptors (mGluR1/5), and subunit-selective NMDARs.

**Results:** Consistent with ionotropic actions, PCP discoordinated CA1 ensemble action potential discharge. PCP caused hyperactivity, and impaired active place avoidance, despite learning the task before PCP. Consistent with metabotropic actions, PCP exaggerated protein-synthesis dependent DHPG-induced mGluR1/5-stimulated long-term synaptic depression (LTD). Pretreatment with anisomycin or the mGluR1/5-antagonist MPEP, both of which repress translation, prevented the PCP-induced discoordination, and the cognitive and sensorimotor impairments. PCP as well as the NR2A-containing NMDAR-antagonist NVP-AAM077 unbalanced translation that engages the AKT, mTOR and 4EBP1 translation machinery and increased protein synthesis, whereas the NR2B-containing antagonist Ro25-6981 did not.

**Conclusions:** PCP dysregulates translation, acting through NR2A-containing NMDAR subtypes, recruiting mGluR1/5 signaling pathways, leading to the neural discoordination that is central to the cognitive and sensorimotor impairments.

## INTRODUCTION

Phencyclidine (PCP) distorts cognition, causes schizophrenia-like symptoms in healthy people and exacerbates symptoms in schizophrenia patients (Cohen et al., 1962; Itil et al., 1967; Abi-Saab et al., 1998). PCP is one of the most dangerous hallucinogens, is increasingly used to lace cannabis and other drugs, and emergency department prevalence of PCP intoxication has been rising (Bush, 2013; Livne et al., 2022). Using the rat hippocampus, spatial cognition, and place cells as a model experimental system, we previously reported that PCP impairs cognition and discoordinates neural activity within and between brain networks that are central to cognitive control (Kao et al., 2017). It is the ability to judiciously process information for attaining goals, and is disrupted in psychotic dysfunction. PCP intoxication that was sufficient to impair cognitive control in a well-learned active place avoidance task, did not alter the basic and spatial tuning discharge properties of single hippocampus place cells, as predicted by the discoordination hypothesis that psychosis-related cognitive impairment results from temporal coordination abnormalities amongst individually normal neural responses (Tononi and Edelman, 2000; Phillips and Silverstein, 2003; Olypher et al., 2006; Uhlhaas and Singer, 2006; Fenton, 2008; Fenton, 2015). Also as predicted by the discoordination hypothesis, PCP discoordinated the sub-second temporal organization of place cell discharge relationships (Kao et al., 2017), consistent with discoordinated activity recorded from neocortex in other psychosis-related manipulations (Hamm et al., 2017). PCP caused theta-modulated medium-frequency gamma oscillations to dominate slow gamma in hippocampus CA1 local field potentials (Kao et al., 2017), biasing neural information processing towards currently perceived information, and away from stored, familiar representations of the environment (Dvorak et al., 2018; Dvorak et al., 2021; Fernandez-Ruiz et al., 2021). The PCP-induced discoordination activated unfamiliar and previously rare hippocampus place cell ensemble discharge representations of a familiar environment, resembling theoretical accounts of delusion and hallucination (Hoffman, 1997), as predicted by attractor models of cognitive discoordination (Olypher et al., 2006; Kao et al., 2017). How might PCP produce these cognition distorting effects, and can they be attenuated?

PCP is a potent uncompetitive antagonist of the glutamatergic N-methyl-D-aspartate receptor (NMDAR) (Lodge and Anis, 1982; Anis et al., 1983) with either direct or indirect dopamine-mimetic effects at dopamine D2 receptors (Adams and Moghaddam, 1998; Seeman et al., 2005; Sershen et al., 2008), but the mechanisms by which PCP produces cognitive and behavioral effects are unknown. Based on the findings that PCP rapidly discoordinates neural discharge, one might reasonably imagine that the cognitive effects are due to the direct ionotropic action of blocking NMDARs to reduce excitation. It is however unclear whether the cognitive and behavioral effects of PCP arise from hypo- or hyperglutaminergic effects (Anand et al., 2000), NMDAR or secondary non-NMDAR actions. PCP alters inhibitory interneuron function (Benardo, 1995; Grunze et al., 1996; Riordan et al., 2018) suggesting it can be disinhibiting. Indeed, there is substantial evidence of network disinhibition effects of uncompetitive NMDAR antagonists including activation by PCP and the psychotomimetic NMDAR antagonists, dizocilpine (MK801) and ketamine (Moghaddam et al., 1997; Hamm et al., 2017). These drugs increase activity of prefrontal (Breier et al., 1997; Vollenweider et al., 1997) and cingulate cortical areas (Holcomb et al., 2005; Rowland et al., 2005) in human neuroimaging studies. In rats, MK801 decreases interneuron firing and increases pyramidal cell firing in prefrontal areas (Homayoun and Moghaddam, 2007) and disrupts the coupling of spiking to local gamma oscillations in the local field potential (Wood et al., 2012). Both ketamine and PCP increase extracellular glutamate in the nucleus accumbens and prefrontal cortex (Liu and Moghaddam, 1995; Moghaddam et al., 1997; Adams and Moghaddam, 1998). Interestingly, the PCP-induced glutamate release in prefrontal cortex is reduced by activation of group II metabotropic glutamate receptors (mGluR2/3) that regulate presynaptic glutamate release (Moghaddam and Adams, 1998). Thus the rapid, schizophrenia-related electrophysiological and behavioral effects of uncompetitive NMDAR antagonists appear to involve an elusive interaction between the NMDAR and mGluR glutamate receptor classes that remains to be clarified providing for the possibility that metabolic effects of PCP administration can also account for the cognitive behavioral effects of the drug.

The present study investigated how PCP might cause neural and cognitive discoordination. Ketamine, which is also an uncompetitive NMDAR antagonist, increases protein synthesis (Li et al., 2010) and like PCP, transiently increases brain-derived neurotrophic factor (BDNF; Takahashi et al., 2006; Autry et al., 2011), which rapidly stimulates protein synthesis (Jourdi et al., 2009), suggesting there could be a role for dysregulated translation. Finally, we noticed similarities between PCP-induced enhancement of gamma oscillations in the hippocampus LFP (Ma and Leung, 2000; Kao et al., 2017) and enhancement of gamma in mice that lack the protein synthesis repressor BC1 RNA when mGluR-stimulated protein synthesis was increased (Zhong et al., 2009). We therefore formulated the working “metabolic hypothesis” that the neural and cognitive discoordination effects caused by PCP arise from excessive protein synthesis.

## RESULTS

### PCP-induced neural discoordination is blocked by the protein synthesis inhibitor anisomycin

The discharge coordination within an ensemble of neurons is estimated by the distribution of correlations between the spike trains of pairs of cells, which in hippocampus CA1 are cell-pair specific and stable (Olypher et al., 2006; Kao et al., 2017). We previously reported that in freely-behaving rats, PCP discoordinates CA1 network discharge, increasing discharge correlations, selectively amongst weakly-correlated cell pairs (Kao et al., 2017). The working hypothesis that the neural and behavioral discoordination effects of PCP are caused by dysregulated translation predicts that the discoordinating effects of PCP will be blocked by pretreatment with a protein synthesis inhibitor such as anisomycin (Fig. 1A).

**Figure 1.**
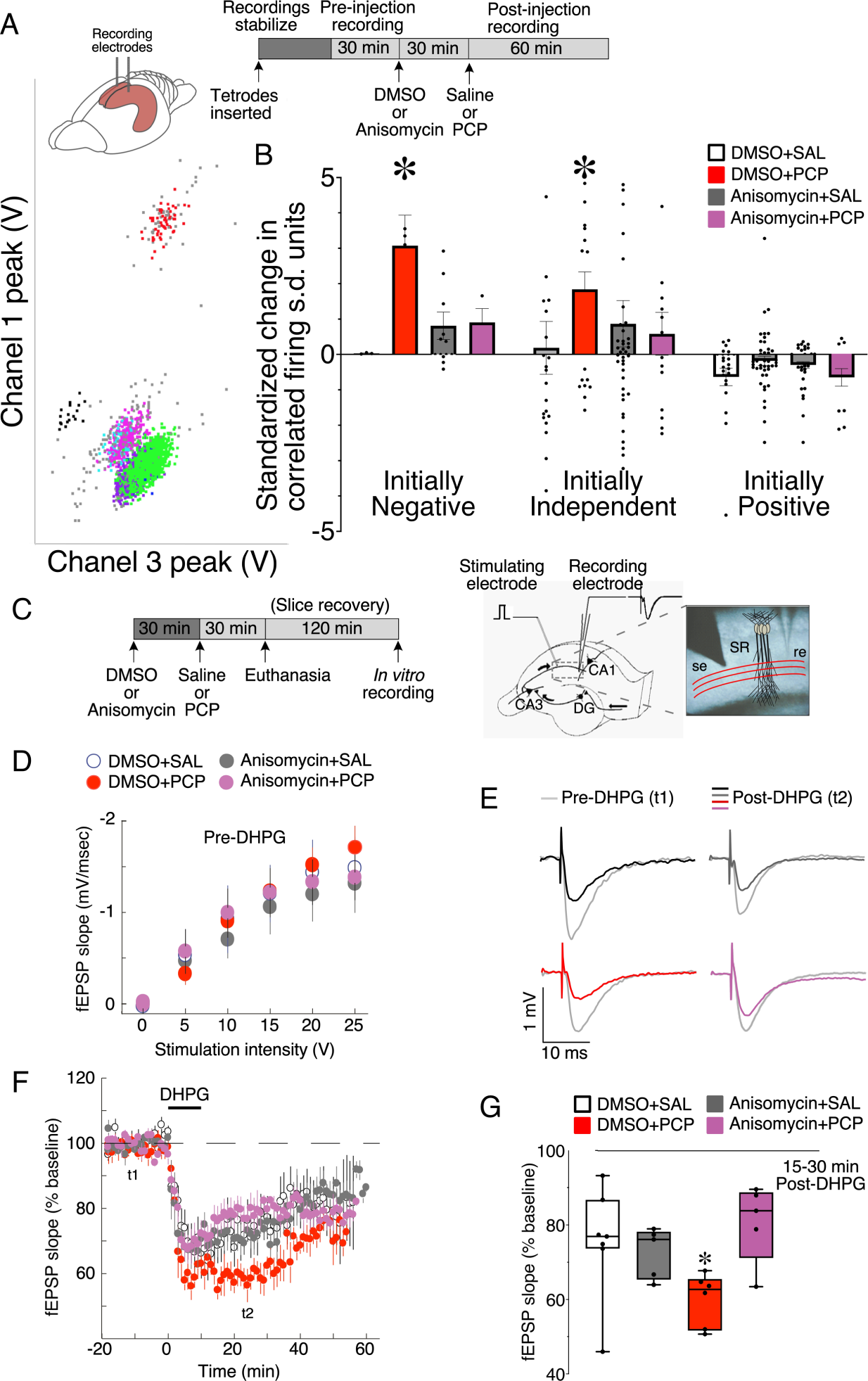
Neural discoordination caused by PCP involves dysregulation of translation. (A) Schematic (top right) illustrating the experimental design. CA1 action potentials were recorded from urethane anesthetized rats using tetrodes, and as shown here, isolated into color-coded single-unit event clusters within the waveform parameter space. (B) The change in Kendall’s correlation (τ) from before to after PCP treatment was computed for all pairs of simultaneously recorded pyramidal cells. Cofiring is computed during 30-min pre- and post-injection at 250-ms resolution. The cofiring changes are grouped by whether cofiring was significantly negative or positive, or independent during pre-injection. PCP (8mg/kg) selectively increases cofiring of initially negatively or independently cofiring cell pairs. This PCP-induced discoordination is blocked by prior administration of (60 mg/kg) anisomysin (Cell numbers: DMSO+SAL n = 57, DMSO+PCP n = 84, Anisomysin+SAL n = 91, Anisomysin+PCP n = 31). (C) Schematic illustrating the experimental design for fEPSP recordings from CA1 *str. radiatum.* Mice received systemic injections before euthanasia and preparation of *ex vivo* hippocampal slices. The drawing and microphotograph illustrate stimulating (se) and recording (re) electrode locations. (D) Prior to DHPG the drugs did not affect evoked fEPSP amplitudes (DMSO+SAL n = 7, DMSO+PCP n = 6, Anisomysin+SAL n = 5, Anisomysin+PCP n = 6). (E-G) PCP-enhanced group I mGluR-stimulated, protein synthesis-dependent synaptic plasticity, measured by 50 µM DHPG-stimulated LTD. Anisomycin pretreatment prevented the mGluR-LTD enhancement by PCP. E) fEPSP examples pre- (t1 = 10 min) and post-DHPG (t2 = 25 min). (F) The time course of the average change in fEPSP slope. (G) The average change between 15 and 30 min after DHPG. *p < 0.03 relative to DMSO+SAL.

Systemic administration of anisomycin (60 mg/kg i.p.) did not change single unit discharge or neural coordination, but pretreatment with anisomycin blocked the 5 mg/kg PCP-induced neural discoordination that was selective to initially negatively or independently cofiring cell pairs, but not positively correlated cell pairs (Fig. 1B). The effects of anisomycin pretreatment, PCP treatment and baseline cell pair category were evaluated by 3-way ANOVA. The effects of pretreatment cell pair category (F_2,241_ = 7.60, p = 6.3x10^-4^) and the interaction between anisomycin and PCP treatment (F_1,241_ = 4.11, p = 0.04) were significant. Post-hoc tests indicated that only PCP alone increased cofiring. These data replicate prior observations in the freely-behaving rat and extend them to an anesthetized condition where PCP-induced hyperactivity cannot contribute to the PCP- induced changes in cofiring. Importantly, these data are the first indication that blocking protein synthesis might be sufficient to prevent the neural discoordination caused by PCP, and therefore that dysregulated protein synthesis could be a mechanism by which PCP causes neural discoordination. We note that anisomycin administration is documented to cause effects other than inhibition of protein synthesis (reviewed by Gold, 2017), which include activation of the MAPK pathway (Shifrin and Anderson, 1999), increased release of a variety of neurotransmitters (Qi and Gold, 2009), and even the silencing of neurons (Sharma et al., 2012). Although control experiments demonstrate that our administration of anisomycin does not silence neurons, we nonetheless point out that the effects of anisomycin cannot only be attributed to protein synthesis inhibition.

### PCP potentiates protein synthesis-dependent group I mGluR-stimulated LTD

We sought additional evidence to more directly test whether PCP-dependent changes of neurophysiology are mediated by dysregulation of protein synthesis. DHPG, the group I mGluR agonist produces robust, protein-synthesis dependent long-term depression (LTD) of synaptic transmission at the Schaffer collateral CA1 synapses in hippocampal slices (Huber et al., 2000; Huber et al., 2001), which is exaggerated by loss of the negative regulators of translation FMRP (Fragile X Messenger Ribonucleoprotein; Darnell et al., 2011) or BC1 RNA (Chung et al., 2017). If PCP dysregulates synaptic plasticity in a similar way, then the *in vivo* PCP treatment should also enhance mGluR-stimulated LTD. Mice were pretreated with anisomycin (60 mg/kg i.p.) or vehicle (DMSO) 30 min before PCP (8 mg/kg i.p.) or saline injections and sacrificed 30 min later (Fig. 1C). Hippocampal *ex vivo* slices were then prepared and treated with 50 µM DHPG to elicit mGluR-stimulated LTD at Schaffer collateral synapses. Before DHPG stimulation, neither treatment with anisomycin nor DMSO affected fEPSP responses (Fig. 1D), but PCP enhanced mGluR-stimulated LTD (Fig. 1E-G). The PCP enhancement measured *ex vivo* was blocked by anisomycin pretreatment, *in vivo*. The interaction between anisomycin and PCP was significant (F_1,19_ = 6.75, p = 0.02). Dunnett’s test showed that only the group treated with the combination of DMSO and PCP was significantly different from the DMSO+SAL control group (p = 0.03, Fig. 1E- G). Despite the protein synthesis dependence of DHPG-induced LTD (Huber et al., 2000; Huber et al., 2001), the anisomycin+SAL group showed strong LTD, indicating again that the effects of systemically-administered anisomycin are complex, certainly when assayed in hippocampal slices more than 3-h after administration to the mouse. There is also evidence of different early and late effects of mGluR1-mediated signaling, which loses its ability to stimulate the neuroprotective, LTD-like mechanisms that require activation of the PI3K-Akt-mTOR signaling pathway (Xu et al., 2007). Nonetheless, these findings provide additional evidence that motivate investigating whether PCP may stimulate translation and reveal an interaction with group I mGluRs.

### PCP changes gamma power and the representational geometry of CA1 population discharge

PCP-induced changes in CA1 discharge population dynamics (Fig. 1B) may be accompanied by changes in local field potential (LFP) oscillations and should alter the representational geometry under which network discharge organizes, perhaps limiting discharge possibilities. Accordingly, we examined PCP’s effect on LFPs and the representational geometry of concurrent CA1 discharge and based on the enhancing effects of PCP on DHPG-induced LTD (Fig. 1G), whether the change is blocked by the group 1 mGluR5 antagonist MPEP. CA1 LFPs and ensemble discharge were recorded from head-fixed mice running in place on a wheel (Fig. 2A,B). LFP spectral power was not altered by pretreatment with saline, DMSO, or MPEP as assessed in the 4 – 8 Hz theta band (F_2,15_ = 0.13, p = 0.9) and the 30 – 100 Hz gamma band (F_2,15_ = 0.43, p = 0.7). Subsequent treatment with PCP did not alter theta compared to saline (F_2,15_ = 1.02, p = 0.4), but PCP increased gamma power and MPEP pretreatment exaggerated the effect (F_2,15_ = 9.01, p = 0.003).

**Figure 2.**
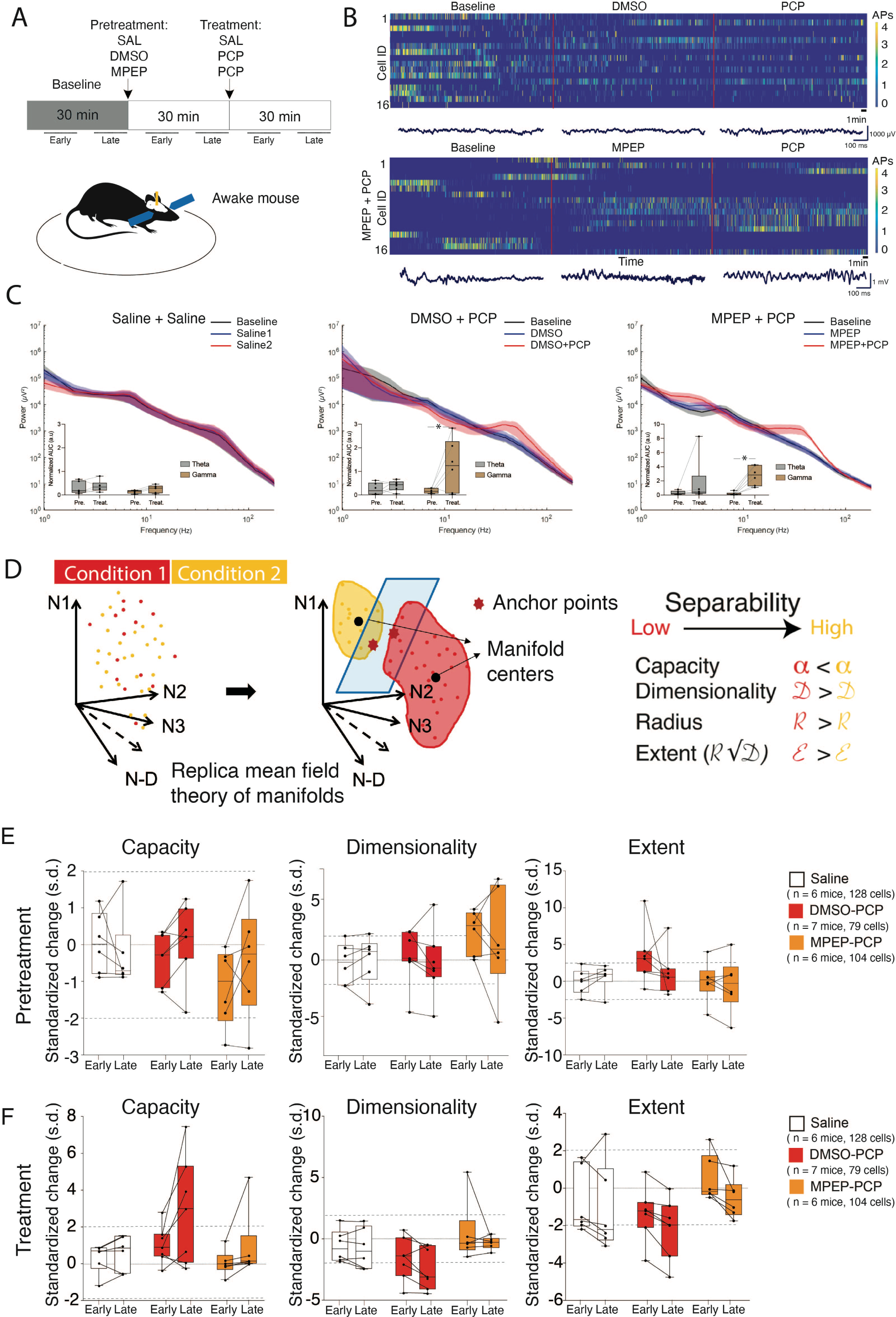
PCP changes the representational geometry of neural population discharge. A) Experimental design. Early: 5-15 min; Late: 20-30 min after SAL/PCP injection. B) Example recordings of CA1 single unit ensembles and LFP across the baseline, pretreatment, and treatment 30-min epochs. C) Average power spectra during baseline, pretreatment, and treatment, with insets that compare the spectral power changes due to pretreatment and treatment on theta and gamma power relative to baseline. Asterisks indicate significant pretreatment vs. treatment differences. D) Cartoon depicting the replica mean field theory used to estimate measures of representational geometry in population dynamics. Measures of three geometric properties E) after pretreatment with saline, DMSO, or MPEP, and F) after subsequent treatment with saline, or PCP indicate that ensemble discharge is unchanged by pretreatment, but becomes more compact after PCP, but the change is blocked by MPEP pretreatment. Standardized changes are measured in s.d. units computed from the fifteen 2-min samples prior to the Pretreatment or Treatment manipulations. Dashed lines indicate 2 s.d. estimates of significant differences caused by pretreatment or treatment.

CA1 ensemble discharge was investigated using replica mean field theory (Chung et al., 2018), within which we computed measures of representational geometry that characterize the linear separability of manifold organization of the population activity temporal dynamics (Fig. 2D). Pretreatment with saline, DMSO, or MPEP could not be distinguished. The two-way Treatment x Time ANOVA confirmed no significant effects of Treatment (F_2,32_’s ≤ 2.53, p’s > 0.1) on the replica manifold measures (Capacity, Extent and Dimensionality). Neither were the effects of Time (F_1,32_’s ≤ 0.71, p’s > 0.4) and the Treatment x Time interactions significant (F_2,32_’s ≤ 0.82, p’s > 0.5). Consistent with PCP increasing correlated activity (Fig. 1B), PCP increased the compactness of the population discharge, measured by computing capacity. dimensionality, and extent of linear separability. By inspection of Fig. 2F, representational capacity appeared to increase over 2 s.d. (p < 0.05) 20-30 min but not 5-15 min post-PCP injection, whereas MPEP pretreatment (70 mg/kg i.p.) prevented these PCP-induced changes. Two-way Treatment x Time ANOVA confirmed significant effects of the Treatment on Capacity (F_2,32_ = 3.73, p = 0.03, PCP > SAL = MPEP+PCP), Dimensionality (F_2,32_ = 5.10, p = 0.01, PCP < SAL = MPEP+PCP) and Extent (F_2,32_ = 3.70, p = 0.04, PCP < MPEP+PCP). However, effects of Time (F_1,32_’s < 3.5, p’s > 0.07) and the Treatment x Time interactions (F_2,32_’s < 0.79, p’s > 0.5) were not significant. These findings in behaving mice confirm that group 1 mGluR antagonism can prevent PCP- induced discoordination.

### The post-training impairment of active place avoidance by PCP depends upon group I mGluR-stimulated protein synthesis

We then investigated in mice, whether PCP impairs an active place avoidance when the drug is administered days after learning the task, like PCP impairs rats doing the task even after intrahippocampal administration that does not cause locomotor hyperactivity (Kao et al., 2017). We also examined whether the impairment also requires protein synthesis as predicted by the working hypothesis (Fig. 3A). Retention of well-learned Room+Arena- active place avoidance (Fig. 3B) that requires cognitive control (Talbot et al., 2018; van Dijk and Fenton, 2018; Chung et al., 2021) was tested 30 min after systemic administration of vehicle or PCP (8 mg/kg). PCP induced hyperactivity (Fig. 3C), and it was prevented by both the anisomycin as well as the MPEP pretreatments. The main effects were significant as well as the interaction (F_2,30_ = 6.24, p = 0.005) and post-hoc tests confirmed that the group that received only PCP moved more than the others. PCP did not abolish place avoidance; mice that received DMSO followed by PCP spent 30.1 ± 10.0 s in the location of the shock zone, which is less than chance (100 s), but more than the other groups (averages < 8 s). This difference could be due to multiple factors, including greater extinction learning under PCP. Nonetheless, the PCP-induced differences are also due to impaired memory, at least in part, and also perhaps impaired sensorimotor integration and cognitive control. This is because the PCP treated mice walked shorter distances before first entering the shock zone, i.e. before they could learn whether or not the shock was still active (Fig. 3D). The impairment was attenuated by pretreatment with anisomycin (60 mg/kg) as well as MPEP (70 mg/kg; Fig. 3D). The 2-way ANOVA on the factors of Pretreatment (vehicle, anisomycin, or MPEP) and Treatment (PCP or vehicle) showed significant main effects as well as the interaction (F_2,30_ = 5.95, p = 0.007). Post-hoc tests confirmed that the group that received PCP without either anisomycin or MPEP differed from all the others. Similar results were obtained when place avoidance memory was measured by the time to first enter the shock zone, which may be more sensitive to the hyperactivity (data not shown; two-way ANOVA interaction: F_2,30_ = 7.23, p = 0.002). These findings demonstrate that in addition to neural discoordination, the PCP- induced cognitive and sensorimotor impairments depend on an anisomycin- and MPEP-sensitive process, which is most likely protein synthesis.

**Figure 3.**
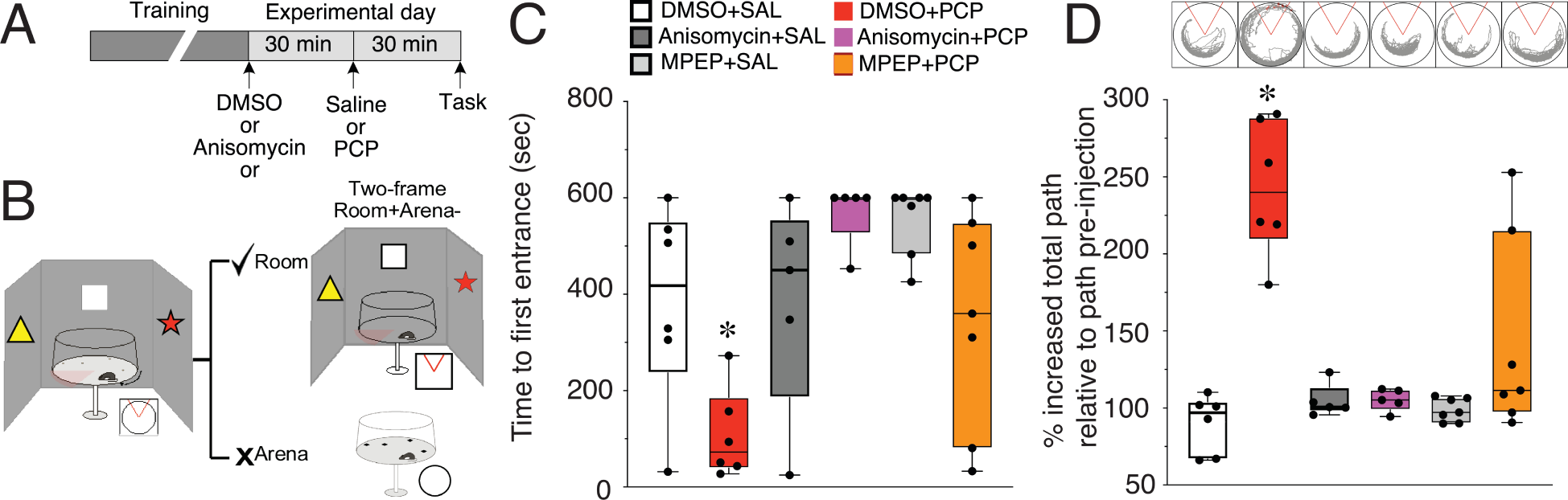
PCP impairs familiar active place avoidance cognitive behavior by dysregulating translation. (A) Schematic illustrating the experimental protocol to test whether pretreatment with anisomycin or MPEP blocks the PCP impairment of familiar active place avoidance (DMSO+SAL n = 6, DMSO+PCP n = 6, MPEP+SAL n = 7, MPEP+PCP n = 7, Anisomysin+SAL n = 5, Anisomysin+PCP n = 6). (B) Schematic illustrating the task concept and environment. (C) Both anisomycin and MPEP pretreatments reduce PCP-induced hyperactivity. D) PCP impairs a well-learned active place avoidance in mice. The impairment is prevented by anisomycin or MPEP pretreatment. Inset: path during test trial for representative mice. *p < 0.05 relative to all groups.

### PCP stimulates group I mGluR-dependent translation

Finally, to directly test whether PCP activates protein synthesis, we used western blot analysis to evaluate the effects of PCP (10 µM), DHPG (100 µM), and the PCP and DHPG combination on activation of the molecular machinery of translation in acute hippocampal slices, compared to vehicle control treatment (Fig. 4A,B). Because the translation machinery represented by ERK1/2 (ERK), mTOR, AKT, and 4EBP1 is active when these proteins are phosphorylated, we measured the ratio of phosphorylated protein/total protein as an indicator of increased activity of these regulatory proteins and more downstream protein synthesis (Topisirovic and Sonenberg, 2011). ARC expression was measured as a downstream indicator of increased translation. Arc is an immediate early gene whose expression is correlated with neural activity (Lyford et al., 1995; Steward and Worley, 2001). One-way ANOVAs followed by Tukey post-hoc tests when appropriate, were performed to compare the drug effects on each molecule. One-way ANOVAs showed significant effects of the treatments on the phosphorylation of ERK, AKT, mTOR, and 4EBP1 as well as on ARC total protein levels (F_3,20_’s ≥ 15.3, p’s < 10^-5^; Fig. 4C,D). Post-hoc tests showed that compared to control, DHPG treatment significantly increased phosphorylation of the translation machinery proteins, as expected. Similarly, PCP enhanced the phosphorylation of translation machinery proteins that reached significance for AKT, mTOR, and 4EBP1 and was marginal for ERK (one-tailed p = 0.05). Combined PCP and DHPG treatment further upregulated the phosphorylation of the same molecules beyond the phosphorylation levels observed with single PCP or DHPG treatments (p’s < 0.01). Complimenting activation of the translation machinery, ARC protein levels were significantly upregulated by individual PCP (p < 10^-3^) and DHPG (p < 10^-3^) treatments compared to the vehicle control and the combined treatment of PCP and DHPG resulted in further increased ARC protein levels beyond the levels observed with single treatments (p’s < 10^-3^). These data directly confirm that PCP interacts with group I mGluRs to promote translation.

**Figure 4.**
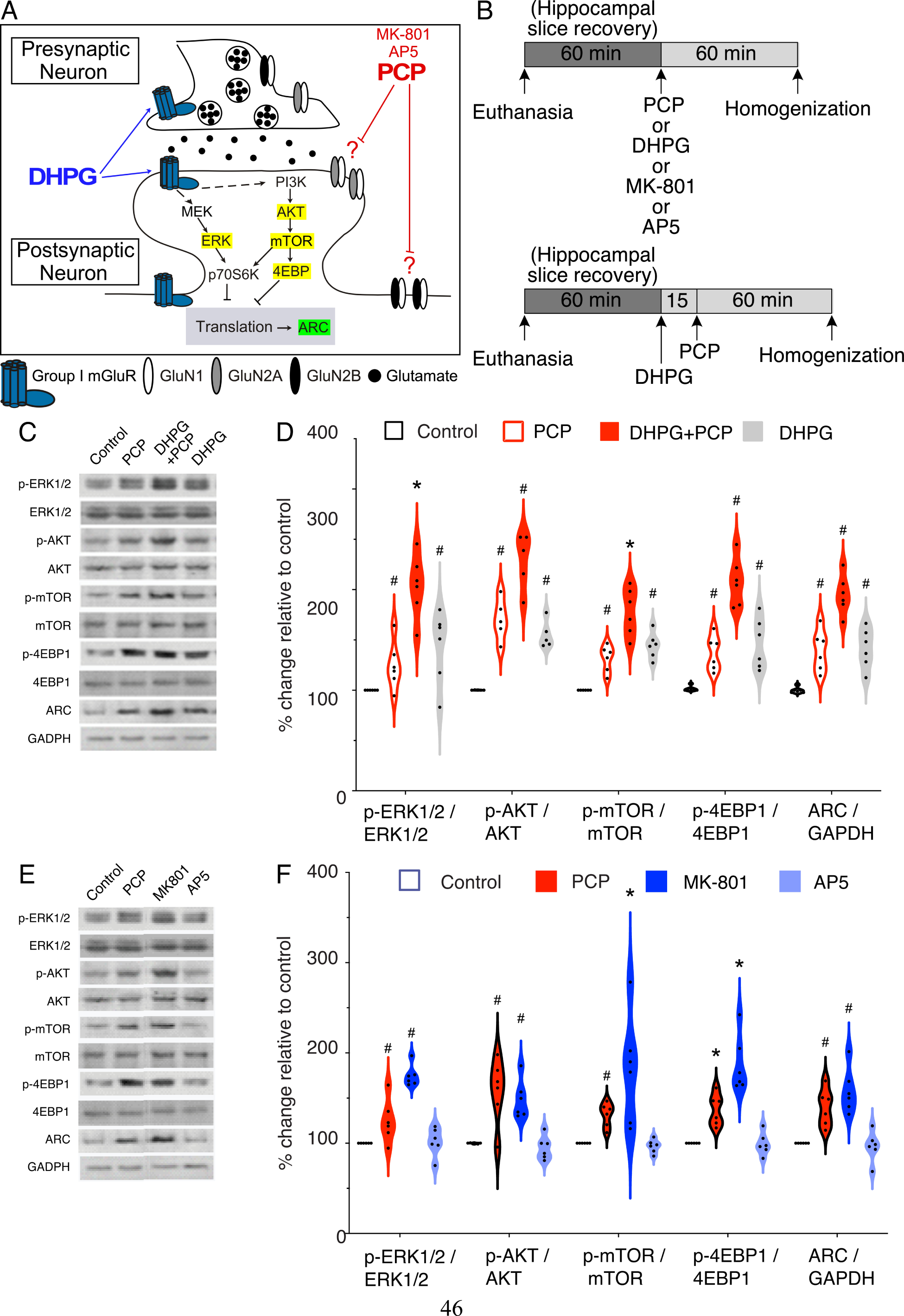
PCP and MK-801 as well as group I mGluR stimulation activates translation, whereas AP5 does not, as measured by expression of the translation machinery components ERK1/2, AKT, mTOR, 4EBP1 and ARC protein. (A) Schematic of known and unknown interactions amongst translation molecular machinery at a CA1 glutaminergic synapse. (B) Schematic experimental design. (C) The *in vitro* effects of PCP on translation machinery in hippocampus were measured by western blot. The n’s = 6 per group. (D) Violin plots show PCP increased activation of translation machinery proteins. (E) Western blot analysis comparing treatment by PCP, MK801and AP5. (F) Translation was stimulated by PCP and MK801 but not AP5. Error bars: ± S.E.M.; Significant effects were confirmed by one-way ANOVA; ^#^p < 0.01 relative to control group, *p < 0.01 relative to all groups.

We replicated the PCP findings in a separate experiment while also testing whether MK801, another uncompetitive NMDAR antagonist, and AP5, a competitive NMDAR antagonist, also activate the translation machinery and upregulate ARC protein. The drug effects were significant in all of the assays (F_4,20_’s <u>≥</u> 9.85, p’s ≤ 10^-4^). Like PCP, relative to vehicle, MK801 (25 µM) upregulated the phosphorylation of mTOR, ERK, AKT, and 4EBP1 as well as increased ARC protein levels (p’s ≤ 0.001; Fig. 4E,F). However, AP5 (50 µM) did not significantly alter the phosphorylation of the analyzed molecules or ARC protein relative to vehicle (p’s ≥ 0.7; Fig. 4E,F). We conclude that PCP deregulates translation, like another uncompetitive NMDAR antagonist, MK801, but unlike the competitive NMDAR antagonist AP5.

### The NR2A-selective antagonist NVP-AAM077 mimics the PCP-induced activation of translation

PCP stimulates translation, but it is unclear how because PCP is an uncompetitive NMDAR antagonist. We next examined if PCP might predominantly act by blocking NR2A- or NR2B- containing NMDA receptors by testing whether antagonists of NR2A- or NR2B-containing receptors mimic the effects of PCP on translation. Acute hippocampus slices were treated with either PCP (10 μM), the NR2A-selective blocker NVP-AAM077 (Romero-Hernandez and Furukawa, 2017), or the NR2B-selective blocker Ro25-6981 (Fischer et al., 1997), each of the blockers at 0.5 μM concentrations.

Compared to vehicle controls, PCP and NVP-AAM077 caused significant increases in the ratio of phosphorylated protein/total protein for ERK, mTOR, AKT, and 4EBP1 (n’s 6-8, F_3,28_’s > 5.76, p’s ≤ 0.003; Fig. 5A). The increased ratios did not differ between PCP and NVP-AAM077 (p’s ≥ 0.3). In contrast, compared to vehicle, the NR2B-selective antagonist Ro25-6981 did not have any significant effects on the above-mentioned proteins (p’s ≥ 0.4). The total protein levels of ERK, mTOR, AKT, and 4EBP1 were not altered by any of the *in vitro* treatments (data not shown). PCP and NVP-AAM077 each significantly increased ARC expression compared to vehicle treatment (p’s ≤ 0.003) whereas Ro 25-6981 did not (p = 0.4; Fig. 5A).

**Figure 5.**
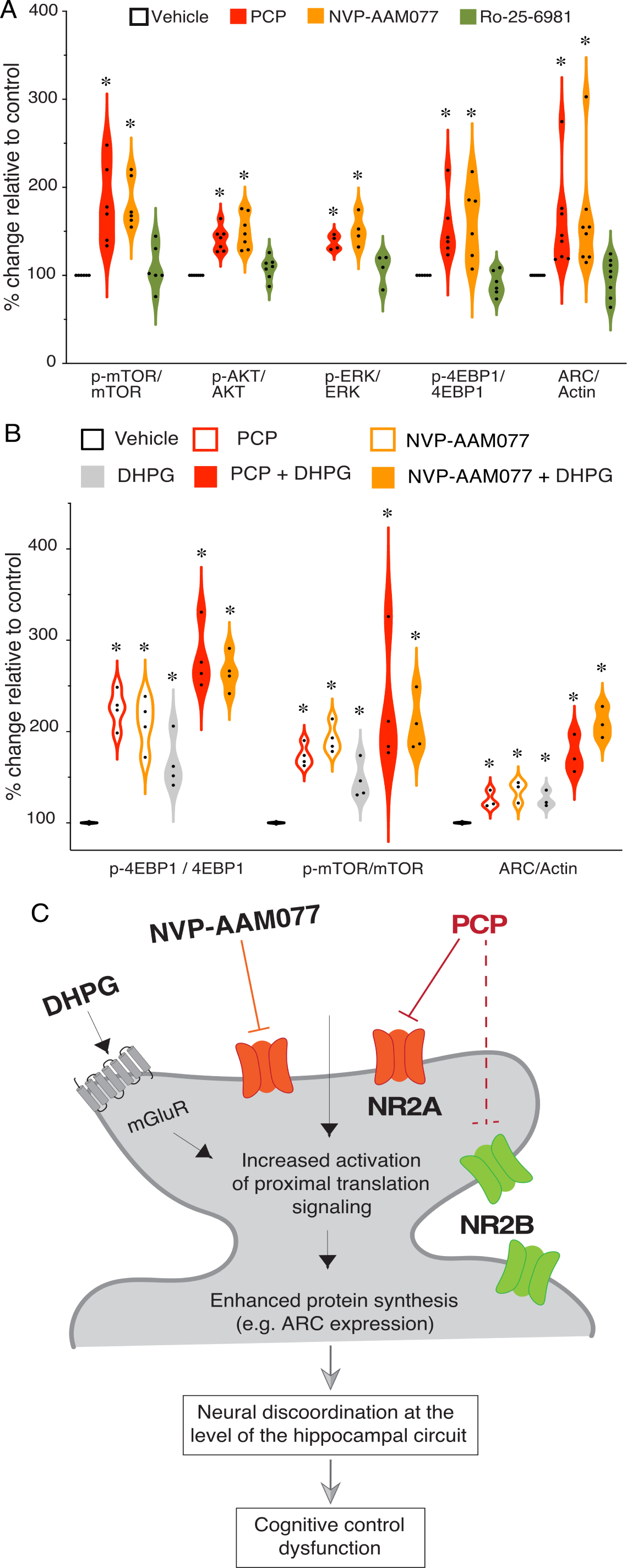
DHPG exacerbates effects of both PCP and the NR2A-containing NMDAR antagonist NVP-AAM077. A) PCP and NVP-AAM077 but not Ro-25-6981 increase phosphorylation of the translation-relevant proteins relative to their total expression. Similar to PCP+DHPG, co-treatment with NVP-AAM077 and DHPG activates the translation machinery more than single treatments. Westerns were run twice for each experimental sample. C) Scheme summarizing the findings and proposed mechanism of acute PCP intoxication. Similar to NVP-AAM077, PCP predominantly blocks NR2A containing receptors in hippocampal spines, causing increased activation of translation signaling and ARC expression, which is exacerbated by mGluR activity. We suggest that the PCP-induced increase in mGluR-dependent protein synthesis contributes to the neural discoordination observed at the level of the hippocampus circuit and the cognitive control dysfunction, measured using the active place avoidance paradigm.

### DHPG exacerbates molecular effects of PCP and NVP-AAM077

Because NMDARs and group 1 mGluRs colocalize (Lujan et al., 1996; Lopez-Bendito et al., 2002), we next tested whether PCP and NVP-AAM077 both act through the group 1 mGluR signaling pathway to stimulate translation. We compared the effect of the two NMDAR antagonists on translation in the presence of the group 1 mGluR agonist, DHPG (100 µM), that we had confirmed combines with PCP to enhance mGluR-stimulated synaptic plasticity that requires translation (Fig. 1D-G). The effects of the Antagonists were all significant (F_2,12_’s > 14.3, p’s ≤ 0.0002; PCP = NVP-AAM077 > Vehicle) as was the effect of DHPG (F_2,12_’s > 7.6, p’s ≤ 0.01). The Antagonist x DHPG interaction was only significant for the effect on ARC (F_2,12_’s > 5.2, p = 0.02; PCP+DHPG = NVP-AAM077+DHPG > PCP = NVP-AAM077 > Vehicle). In summary, DHPG treatment alone increases the phosphorylation of 4EBP1 and mTOR and enhances ARC expression, as expected (Fig. 5B), whereas co-treatment of PCP or NVP- AAM077 with DHPG further enhances translation signaling.

## DISCUSSION

### Main findings

Being aware of off-target pharmacological effects, we used multiple assays at distinct levels of biological organization as well as agonist and antagonist manipulations to discover that PCP disrupts neural coordination and cognitive control by dysregulating translation. We find that PCP discoordinates the temporal organization of spiking in urethane-anesthetized rats by coactivating cell pairs that were unlikely to discharge together before PCP (Fig. 1). PCP also increases CA1 gamma oscillations (Fig. 2C; Kao et al., 2017) and discoordinates CA1 neural population discharge from the awake mouse CA1, resulting in more stereotyped and distinctive neural population dynamics (Fig. 2). These findings replicate and extends previous findings in freely-behaving animals, since PCP-induced hyperlocomotion and other behavioral confounds cannot account for the present electrophysiological findings (Kao et al., 2017). This specific form of PCP- induced neural discoordination is similar to the discoordination effects of MK-801 (Szczurowska et al., 2018) as well as the effects of hippocampal disinhibition by TTX inactivation of the contralateral hippocampus (Olypher et al., 2006). Second, we report that PCP also impairs conditioned place behavior in a familiar active place avoidance task that requires cognitive control demonstrated behaviorally (Fig. 3), as well as with electrophysiology and calcium imaging concurrent with behavior (Talbot et al., 2018; van Dijk and Fenton, 2018; Chung et al., 2021). This extends to mice a previous observation in rats (Kao et al., 2017). Third, we demonstrate that anisomycin and MPEP pretreatments prevent the electrophysiological, cognitive behavioral, and the sensorimotor effects of PCP. We also demonstrate that PCP enhances group 1 mGluR-stimulated LTD, a protein synthesis-dependent type of synaptic plasticity, whereas PCP itself does not (Fig. 1). These findings argue that the cognition-impairing effects of PCP arise from dysregulated group I mGluR-stimulated protein synthesis. We then demonstrate that PCP increases proximal, translation-signaling in both rat (Fig. 4) and mouse (Fig. 5) hippocampus as indicated by enhanced ARC expression and increased phosphorylation of the following proteins: ERK, AKT, mTOR, and 4EBP1. This is consistent with the data showing that treatment with either anisomycin or MPEP can prevent PCP-induced forms of neural discoordination (Figs. 1,2). Finally, by mimicry experiments, we demonstrate that the PCP effects on proximal activation of translation can be the result of PCP’s antagonist action on NR2A-containing NMDARs leading to group 1 mGluR stimulation (Figs. 4,5).

### Comparison with neocortical effects of NMDA receptor antagonists: The discoordination hypothesis

It is remarkable that both PCP and MK-801 (another uncompetitive NMDAR antagonist) as well as contralateral hippocampal inactivation with TTX all cause the same specific pattern of neural discoordination as well as a specific cognitive behavioral deficit in the <u>use</u> of established spatial memories, rather than in place learning or information storage *per se (Stuchlik and Vales, 2005; Wesierska et al., 2005; Olypher et al., 2006; Kubik et al., 2014; Kao et al., 2017)*. These observations are predicted by the discoordination hypothesis (Fenton, 2008; Fenton, 2015), which asserts that psychosis-related cognitive deficits, such as impaired cognitive control, arise because processing multiple streams of information is corrupted by aberrant coordination of neural activity, in which cells that normally discharge together and cells that normally do not discharge together fail to maintain their appropriate timing relationships, despite maintaining their individual response properties (Tononi and Edelman, 2000; Phillips and Silverstein, 2003; Olypher et al., 2006; Uhlhaas and Singer, 2006; Lee et al., 2012; Lee et al., 2014; O’Reilly et al., 2014). The idea derives directly from cell assembly (Hebb, 1949) and other ensemble hypotheses for how information is represented in the brain, which seem valid in hippocampus (Harris et al., 2003; Itskov et al., 2008; Fenton et al., 2010; Kelemen and Fenton, 2010b; Dragoi and Tonegawa, 2011; Park et al., 2011; Silva et al., 2015; van Dijk and Fenton, 2018). Ensemble hypotheses assert that streams of information are represented by the coordinated temporal spiking relationships amongst cells in distributed representations. It follows that temporal discoordination will derange representations and the judicious use of information, predictions that are confirmed by the present findings.

Indeed, prior work has demonstrated electrophysiological discoordination after PCP and other NMDA receptor antagonists. Like the present work in hippocampus, ketamine and MK-801 induced increased gamma band power (Pinault, 2008) as well as increased correlated discharge in neocortex (Hamm et al., 2017), which in the case of ketamine is accompanied by diverse region-specific increases and decreases in metabolism assessed by 2-deoxyglucose imaging (Dawson et al., 2013) and increased glutamate and dopamine release (Moghaddam et al., 1997). The discoordinating effects of MK-801 on prefrontal cortical neurons is likely due to drug-dependent decreases of inhibitory control and consequent increases of excitatory firing (Jackson et al., 2004; Homayoun and Moghaddam, 2007), whereas we did not see increases in discharge rates in the present or prior study of hippocampus, likely due to especially strong firing rate homeostatic mechanisms in hippocampus (Buzsaki et al., 2002; Kao et al., 2017). While we did not assess whether the effects of PCP were reversible and the spike train discoordination lasts less about a hour (Kao et al., 2017), the metabolic effects are likely to have produced enduring effects, which in the case of attenuated LTP have been described to persist for weeks following a single dose of MK-801 (Manahan-Vaughan et al., 2008). We provide evidence that the metabolic effects of PCP are likely due to antagonism of NR2A-containing NMDA receptors. This does not imply that the electrophysiological effects are also due to NR2A antagonism, although deletion of NMDA receptors in parvalbumin-expressing neurons is sufficient to express enhanced gamma (Carlén et al., 2012). Low doses (≤10 mg/kg) of MPEP are reported to decrease prefrontal cortical neuronal discharge (Homayoun and Moghaddam, 2010), but we did not observe such effects of high dose MPEP in hippocampus CA1, illustrating that the electrophysiological drug effects are likely to be subregion specific and dependent on the region’s particular network properties.

### Mechanism of PCP-induced cognitive impairments

The discoordinating actions of PCP have been observed rapidly, within a minute in the awake rat (Kao et al., 2017) suggesting ionotropic effects, consistent with alteration of interneuron function (Benardo, 1995; Grunze et al., 1996; Homayoun and Moghaddam, 2007), but as the present findings demonstrate, PCP effects are also metabolic, which can also be rapid, within 5 min (Tsokas et al., 2005; Waung et al., 2008). PCP transiently increases BDNF (Takahashi et al., 2006), which rapidly stimulates protein synthesis via the mTOR pathway (Fig. 4; Jourdi et al., 2009). PCP’s impairment of cognition also appears related to dysregulation of mGluR-stimulated translation, because the PCP effects were blocked by either anisomycin or MPEP and are exacerbated by group 1 mGluR stimulation (Fig. 3,4). The PCP effects appear to be through both the mGluR-MEK/ERK and the mGluR-mTOR translation pathways (Fig. 4,5), although this requires further investigation. The related psychotomimetics ketamine and MK801 affect both pathways (Ahn et al., 2006; Yoon et al., 2008; Hong et al., 2010; Li et al., 2010). While controversy persists over which translation pathways are more relevant for ketamine’s effects (Li et al., 2010; Autry et al., 2011; Carrier and Kabbaj, 2013; Gideons et al., 2014), the present findings with AP5, MK-801, Ro25-6981, and NVP-AAM077 demonstrate that different NMDAR antagonists have different effects on translation, and so not unsurprisingly, also different potential for intoxication and therapy.

The present findings indicate that PCP enhances mGluR-dependent translation to produce cognitive impairment. Fig. 5C summarizes the findings, but there are multiple possibilities for how PCP interacts with mGluR-stimulated translation and we note that MPEP itself is a weak NMDAR antagonist (O’Leary et al., 2000). The most parsimonious possibility is that by blocking NR2A- containing NMDARs, PCP makes more glutamate available to stimulate colocalized mGluR’s (Lujan et al., 1996; Lopez-Bendito et al., 2002), but it is not obvious why uncompetitive open channel blockade by PCP should reduce glutamate binding to NMDARs (Ferrer-Montiel et al., 1995; Monaghan and Jane, 2011). Alternatively, PCP can increase glutamate signaling onto principal cells by preferentially blocking NR2A-containing NMDARs on interneurons (Benardo, 1995; Grunze et al., 1996; Homayoun and Moghaddam, 2007) or possibly by interacting with astrocytic gliotransmission that is sensitive to group I mGluR stimulation (Fellin et al., 2004; Fiacco et al., 2007). Aberrantly increased glutamate signaling would increase mGluR stimulation, initiating a molecular cascade that increases translation at synapses causing a dysregulation of excitation and inhibition (Gladding et al., 2009). A weak prediction of this model is increased principal cell firing, which is not observed (Fig. 2B; Kao et al., 2017). However, the observation may be the result of tight coupling between excitation and inhibition and mechanisms of firing rate homeostasis in hippocampus (Buzsaki et al., 2002; Olypher et al., 2006).

Interactions between NMDARs and mGluRs are likely to contribute to the interplay between PCP and mGluR signaling, which is complex, and all the more so because mGluR3 activation potentiates mGluR5 activity (Di Menna et al., 2018). Chemical stimulation of NMDARs enhances repression of translation by micro RNAs (Antoniou et al., 2014). By blocking NR2A-containing NMDARs, PCP would derepress translation by this mechanism. It is also likely that mGluR stimulation will itself derepress translation because DHPG treatment of primary neuronal cultures stimulates presynaptic mGluRs, resulting in AMPA receptor (AMPAR) endocytosis which has been shown to result in enhanced translation (Park et al., 2008; Luscher and Huber, 2010; Xu et al., 2013; Antoniou et al., 2014). These NMDAR and mGluR mechanisms of stimulated translation may account for the present findings that PCP blockade of NMDARs promotes translation, that the effect of mGluR stimulation by DHPG is additive, whereas mGluR5 antagonism attenuated PCP’s electrophysiological (Fig. 2) and behavioral (Fig. 3) effects. PCP also enhanced ARC synthesis, which is implicated in downregulation of surface-bound AMPAR and LTD (Waung et al., 2008; Niere et al., 2012). However, because the effects of PCP intoxication on cognitive and neural representation are transient, lasting ∼ 1h (Kao et al., 2017), persistent changes in synaptic plasticity are unlikely to explain our observations, but may account for the derangements that follow chronic exposure (Linn et al., 1999; Balla et al., 2003; Amitai et al., 2011).

The prevention of PCP-induced hyperlocomotion and active place avoidance impairments by high dose of MPEP (Fig. 3) contradicts reports that group I mGluR antagonists exacerbate the effects of PCP and related NMDAR antagonists (Henry et al., 2002; Kinney et al., 2003; Campbell et al., 2004; Pietraszek et al., 2004; Pietraszek et al., 2005; Doreulee et al., 2009) and reports that positive allosteric modulators have either beneficial or no effects (Kinney et al., 2003; Schlumberger et al., 2010). Most, but not all of these studies assayed effects on sensorimotor gating, whereas we focused on cognitive effects. Nonetheless, assessment differences are insufficient to explain the outcome differences because we also observed that MPEP prevents PCP-induced hyperlocomotion (Fig. 3C). Prior studies administered mGluR5 antagonist doses up to 10 mg/kg, whereas we pretreated the mice with 70 mg/kg MPEP, 30 min before PCP to ensure effectiveness during training despite the relatively rapid metabolism of MPEP in mice (Anderson et al., 2002; Anderson et al., 2003). In pilot work, we had observed that lower doses of MPEP did not attenuate PCP-induced hyperactivity, and that sometimes PCP could transiently cause hyperactivity immediately after it was administered, even after the 70 mg/kg MPEP pretreatment. The present findings are clear and robust at the levels of *ex vivo* and *in vivo electro*physiology and cognitive behavior, demonstrating that mGluR5 antagonists can be procognitive, apparently by dampening translation which PCP can stimulate. These metabolic effects suggest a potential strategy for reducing the effects of PCP intoxication, which appears distinct from the group II mGluR agonist strategies that were once considered for procognitive antipsychotic development (Moghaddam and Adams, 1998; Patil et al., 2007; Weinberger, 2007).

### Procognitive antipsychotic drug development

The PCP-induced abnormalities in cognitive behavior, neural discoordination, and molecular signaling demonstrate there may be utility of the acute PCP animal model for studying the core cognitive symptoms of schizophrenia, which we understand contrasts with the waning interest in PCP in light of prior lack of success, and the availability of more selective antagonist compounds and genetic models of hypothesized etiological causes of schizophrenia. Indeed, acute PCP intoxication has been the most extensively used experimental preparation for antipsychotic drug development, where the focus has been on treating the positive symptoms of psychosis. Given the current interest to develop drugs that target cognitive symptoms in schizophrenia, the present demonstration of a set of PCP-induced cognition related deficits indicates that the use of the PCP model can also be extended to the search for a procognitive antipsychotic. In this regard, it is important to consider that PCP-induced cognitive control deficits can be dissociated from the long-established hyperlocomotion that is thought to model the positive symptoms (Kao et al., 2017). Infusing PCP directly into the hippocampus impaired cognitive control measured by place avoidance without causing hyperlocomotion, so long as the drug increased 60-100 Hz mid-frequency gamma oscillations (Kao et al., 2017). The dissociation could help understand why treatment strategies that were developed by targeting hyperactivity and other sensorimotor abnormalities turned out to be inadequate for the cognitive symptoms (Carter and Barch, 2007). The present work and related work of Kao et al., (2017) also provides compelling evidence that although NMDAR antagonism spares established memory, it can devastate the judicious use of information when sources of cognitive interference abound such as during two-frame avoidance or initial place learning in the water maze (Bannerman et al., 1995; Saucier and Cain, 1995). These findings suggest it may be necessary to investigate cognitive control, in addition to learning and memory *per se*, in research directed at schizophrenia (Gilmour et al., 2013) as well as other fields (Talbot et al., 2018; Chung et al., 2021) as predicted by the discoordination hypothesis (Phillips and Silverstein, 2003; Fenton, 2008; Fenton, 2015). While these behavioral findings on their own suggest that acute PCP intoxication could with the right assays be exploited to model the core cognitive deficits in schizophrenia, the utility of the model lie in the theoretical importance of the findings. The electrophysiological findings provide unambiguous evidence that PCP discoordinates electrical neural activity between neurons (Fig. 1,2) with minimal consequences on the response characteristics of individual cells during normal behavior (Kao et al., 2017) as predicted by the discoordination hypothesis. The findings with anisomycin and MPEP, as well as our direct assays of the translation molecular machinery, indicate that the PCP-induced cognitive impairment is mediated or at least modulated by dysregulated translation, consistent with findings in DISC1 and PKR-like ER kinase (PERK) mutant mice that mimic gene alterations in schizophrenia and implicate excessive translation in the cognitive deficits associated with the disease (Trinh et al., 2012; Zhou et al., 2013). Despite stimulating translation and consequent BDNF synthesis, which has antidepressant effects when elicited by ketamine, another uncompetitive NMDAR-antagonist (Takahashi et al., 2006; Autry et al., 2011), the effects of PCP are debilitating. This highlights the need for caution and rigor in attempts to develop mechanistically-related ketamine antidepressant therapies. The findings that doses of PCP that impair cognition also discoordinate neural activity suggest that normalizing neural discoordination could be a procognitive therapeutic target. Although this may be accomplished by correcting a proximal cause such as dysregulated translation, procognitive effects can also be achieved by correcting the discoordination itself, using pharmacological treatment to restore excitation-inhibition balance (Lee et al., 2012), or even by harnessing the plasticity that can accompany appropriate cognitive experience (Lee et al., 2014; Pavlowsky et al., 2017; Chung et al., 2021).

## MATERIALS AND METHODS

All behavioral and electrophysiological methods have been published elsewhere (Olypher et al., 2006; Chung et al., 2017; Talbot et al., 2018; Dvorak et al., 2021). Dorsal hippocampus pyramidal cell discharge and evoked-potential population responses were analyzed for Figures 1 and 2, behavior was analyzed for Figure 3, and protein synthesis was analyzed for Figures 4 and 5.

### Drugs

Phencyclidine (PCP, 1-(1-Phenylcyclohexyl)piperidine hydrochloride, pH 7.4; Sigma, St. Louis, MO, Cat. No. P3029) and 2-methyl-6-(phenylethynyl)pyridine (MPEP; Tocris, Ellisville, MO, Cat. No. 1212/10), solutions were made fresh in sterile saline before use. Anisomycin (Sigma, St. Louis, Cat. No. A9789), solution was made fresh in DMSO before use. (S)-3,5-Dihydroxyphenylglycine, (DHPG, Tocris, Ellisville, MO, Cat. No. 0805), D-2-amino-5-phosphonopentanoate (D-AP5, Tocris, Ellisville, MO, Cat. No. 0106 ), and Dizocilpine ((+)-MK801, Sigma-Aldrich, St. Louis, MO, Cat. No. M107), Ro 25-6981 maleate (R&D Systems, Cat.No. 1594), and NVP-AAM077/PEAQX (Sigma, Cat.No. P1999) were dissolved in sterile water.

### Animals

All animals were maintained on a 12:12 light-dark cycle. Adult male Long-Evans rats and C57BL/6 mice were 2-3 months during the experiments. All procedures conformed to institutional and NIH guidelines for the treatment of vertebrate animals and were approved by the SUNY Downstate and NYU Institutional Animal Care and Use Committees.

### *In vivo* electrophysiology – Acute recordings under anesthesia

The *in vivo* methods to record single unit discharge from urethane-anesthetized rats have been described in prior work (Olypher et al., 2006). Briefly, rats were anesthetized with urethane (1.2g/kg i.p.) then mounted in a stereotaxic frame with precision micromanipulators (TSE Systems GmbH, Bad Homburg, Germany). The scalp was resected and crainotomies were performed at AP -3.8 mm, ML ±2.5 mm relative to bregma to provide electrode access to the dorsal hippocampus. The dura was cut, and tetrode-configured electrodes made from 4 twisted 25-μm nichrome wires (impedance 50-200 Ko), were lowered to the recording targets using the stereotaxic micromanipulators, guided by electrophysiological landmarks in the local field potentials and single unit activity. Data collection began once stable single unit ensemble activity could be isolated. A baseline recording was made for at least 30 minutes and then the rat received an i.p. injection of the 5 mg/kg PCP or vehicle solution. Recordings continued at least one hour after the injection.

### Recording setup

Extracellular, 2-ms tetrode action potential waveforms were buffered by a custom pre-amplifier, band-pass filtered (low-cut frequency 300 or 360 Hz, hi-cut frequency 5-10 kHz), amplified (5,000-10,000 times), and digitized (32 or 48 kHz). The digital signals were stored in real-time using either custom software (AcX, A.A. Fenton) or a commercial system (dacqUSB, Axona Ltd., St. Albans U.K).

The neuronal activity was recorded under anesthesia, for at least 30 min in the pre-injection period. Afterwards the rats received a systemic injection of the control, or behaviorally-effective dose of PCP. Post-injection recordings lasted at least 60 min. All data were analyzed off-line. The data recorded in the 30-min pre-injection and the first 30-min of post-injection interval were analyzed, during which PCP drug status was in an effectively steady state (Kao et al., 2017).

### Single unit identification

The action potentials emitted by different cells were discriminated off-line using custom waveform parameter clustering software (Wclust, A.A. Fenton). Each action potential waveform was characterized by parameters that included the positive and negative peak voltages on each tetrode wire, the voltage at user-selected times relative to the spike onset, the waveform energy, principal components, and others. Single units were classified as those waveforms that formed clusters in the waveform parameter space according to quantitative criteria of isolation from all other spikes (Iso_BG_ ≥ 4) and isolation from the most similar cluster of spikes (IsoI_NN_ ≥ 4) (Neymotin et al., 2011).

### *In vivo* electrophysiology – Recordings in head-fixed behaving mice

Mice under isoflurane anesthesia were surgically prepared for head-fixed recordings. A titanium head plate was attached to the mouse’s skull using dental cement and the exposed skull was covered with KwikSil, a low toxicity adhesive (World Precision Instruments, Sarasota, FL) and protected by attaching a plastic cup. All mice were allowed at least 1 week to recover. A secondary surgery was performed immediately before the experiment. The plastic cup and KwikSil were removed, and a craniotomy was made at 1.85 AP, ± 1.20 ML relative to bregma to enable electrode placement targeting dorsal hippocampus. After recordings the KwikSil protective cup assembly was reattached to prevent infection.

Once the mouse was head-fixed by the titanium plate, A Neuronexus (Ann Arbor, MI) silicon probe was positioned to record from CA1 using a Kopf stereotaxic arm. Signals were filtered between 300 Hz and 10 kHz and sampled at 30 kHz for single unit recording and filtered between 0.1 Hz and 1 kHz and digitized at 2 kHz to record the concurrent LFPs.

LFPs at the CA1 recording sites were averaged for analysis. Powerspectra were computed using the pwelch() function in MATLAB. Theta (4 - 8 Hz) and gamma (30-100 Hz) band power were computed, and for comparisons, normalized by the spectra computed from the preceding baseline recording. The normalized spectra were compared across pretreatment with saline, DMSO and MPEP, and compared separately across the saline-saline, DMSO-PCP, and MPEP- PCP treatment groups.

Single units were sorted using a published open-source algorithm Kilosort2 (Pachitariu et al., 2016) that takes advantage of GPU processing to improve algorithm performance. After automated clustering of the data, only units with < 20% estimated contamination rate with spikes from other neurons that were computed from the refractory period violations relative to expected. We also excluded units with non-characteristic or noisy waveforms to identify single units.

### Classification of cell types

Single units were classified as complex-spike or theta cells according to published criteria (Ranck, 1973; Fenton et al., 2008; Kelemen and Fenton, 2010a). Complex-spike cells appear to be pyramidal cells, whereas theta cells are likely local interneurons (Fox and Ranck, 1975). Pyramidal cells were judged to have long-duration waveforms (> 250 µs), low discharge rate (< 2 AP/s) and a tendency to fire in bursts (peak inter-spike interval < 10 ms). Interneurons had short-duration waveforms (< 250 µs), high discharge rate (> 2 AP/s), and did not tend to fire in bursts.

### Functional coupling of cell pairs

The functional coupling of spike trains from pairs of cells was estimated using Kendall’s correlation (Press et al., 1993), which was computed as previously described (Olypher et al., 2006). The correlation was computed between the time series of spike counts from the two cells. The time series was generated by counting the number of spikes the cell fired during each 250-ms interval.

### Geometric properties of high-dimensional neural manifolds

The replica mean field theory of manifolds computes the geometric properties of neuronal population activity without assumptions on the topology or dimensionality of the high dimensional neural data (Chung et al., 2018). The replica-manifold theory assumes that neural manifolds may be defined by D+1 coordinates, such that one coordinate is needed to describe the location of the neural manifold center, and D coordinates to account for the axes that define the manifold variability. Ensemble neural activity organizes on P manifolds sampled M times along N different feature dimensions, i.e., individual cells. In this analysis, there are P manifolds that correspond to P recordings of an animal in different conditions within a recording.

We use three geometric properties to characterize high-dimensional neural data: capacity, dimensionality, and extent. Classification capacity, α_c_ is defined as the maximum number of manifolds that can be linearly separated given a random assignment of binary labels to manifolds. Capacity can be understood as a high-dimensional measure of overlap between manifolds, a measure of manifold compactness. Similar to the concept of support vectors in linearly separable points, in replica theory, the high dimensional separating hyperplane is defined by a linear combination of what we call manifold anchor points. Anchor points uniquely define the separating plane between manifolds. Anchor points depend on the location and orientation of the manifolds as well as the randomly assigned binary labels. For a given manifold, one can define a statistical distribution of anchor points. From the distribution of anchor points, we can learn about two geometric properties of high dimensional manifolds: the effective radius, R_M_, defined as the square root of the total variance of the anchor points, normalized by the average norms of the manifold centroids, and the effective dimension, D_M_ which captures the spread of the anchor points along the different manifold axes. We further define the manifold total extent, a measure of the space occupied by the high dimensional neural manifold as the mean R_M_√(D_M_). According to the replica mean field theory of manifolds, larger D_M_ and R_M_√(D_M_) of the manifolds are linked to statistically lower linear separability between manifolds, and less distinction among themselves.

### *Ex vivo* hippocampal slice electrophysiology

Mice were systemically pretreated with anisomycin (60 mg/kg) or the DMSO vehicle and 30 min later with PCP or saline. Mice were sacrificed 30 min after the second treatment to obtain hippocampal slices (400 μm). Slices were incubated for 2 hours in oxygenated aCSF (in mM: 125 NaCl, 2.5 KCl, 1 MgSO_4_, 2 CaCl_2_, 25 NaHCO_3_, 1.25 NaH_2_PO_4_ and 25 Glucose), and then were placed in a submerged chamber subfused with aCSF at 35-36°C for recording (Pavlowsky and Alarcon, 2012; Chung et al., 2017). A pair of stimulation (bipolar; FHC & Co, ME, USA) and recording electrodes (borosilicate glass pipette filled with aCSF; 5-10 mΩ) was used to evoke and record field excitatory postsynaptic potentials (fEPSP) at the CA1 *stratum radiatum*. Stimulus-response curves were created by delivering square pulses (50 μsec) at increasing voltages (0- 25V). For long-term synaptic depression (LTD) studies, fEPSP responses were set at 40% of the maximum slope amplitude and sampled once per minute. After a stable baseline was established, the group I mGluR agonist DHPG (25 μM) was washed onto the slices for 10 min to induce LTD. LTD slope amplitudes (percentage change from baseline at 15-30 min) between conditions were compared.

### Behavior - Active place avoidance task

#### Setup

An individual mouse was placed on a metal disk-shaped arena (40 cm in diameter). The arena was elevated 76 cm above the floor and centered in 3x4 m^2^ room surrounded by opaque curtains and various items (Fig. 3B). The mouse was constrained to remain on the disk by a 40-cm-high transparent wall. The mouse’s position was tracked every 33 ms from an overhead camera using digital video spot tracking software (Tracker, Bio-Signal Group Corp., Acton, MA).

The two-frame Room+Arena- task variant was used. The arena rotated at 1 rpm and an unmarked 60° shock zone was fixed at stationary room coordinates. The arena floor was made of parallel metal rods, configured as five electrical poles for delivering a mild electrical shock (0.2 mA, 60 Hz, 500 ms) scrambled across the rods. Under software control, shock was delivered when the mouse was in the shock zone for 500 ms. The shock was repeated every 1.5 seconds until the mouse left the shock zone. To avoid a shock the mouse had to use the relevant stationary distal room cues to localize itself and the positions of the shock zone. Because the arena was rotating the mouse also had to ignore the irrelevant information from the rotating spatial frame. Note that this shock regime is no more stressful than walking freely in the arena (Lesburgueres et al., 2016).

#### Behavioral Protocol

The goal was to test the effect of PCP on the ability of mice to do active place avoidance after the task had been learned and the memories established, extending the finding that PCP impairs familiar active place avoidance in rat to the mouse (Kao et al., 2017). The mice were first habituated to the environment during a 10-min trial on the stationary arena with no shock. The shock and arena rotation were then turned on and mice were given three 10-min training trials with 30-min inter-trial intervals for 4 days. On day 5, each mouse received DMSO, MPEP (70 mg/kg), or anisomycin (60 mg/kg) 30 min before either saline or PCP (8 mg/kg). Thirty min after the 2^nd^ treatment, mice were tested in conditions that were identical to the training but without shock.

The time series of the animal’s positions were analyzed offline. The time to first enter the shock zone was measured because it increases as the animals learn to avoid shock and can be interpreted as an index of the ability to retrieve the place avoidance memory. The distance the animals walked was used to estimate locomotion because it can characterize hyperactivity, which is known to be caused by PCP (Patil et al., 2007).

### Protein expression - Western blotting

We used western blot analysis to evaluate the effects of PCP and other drugs on the activation of the protein translation machinery. Specifically, extracellular-signal-regulated kinase (ERK), mammalian target of rapamycin (mTOR), protein kinase B (AKT), and 4E-binding protein (4E- BP1) are involved in proximal translation signaling; ERK is also involved in many other cellular metabolic processes (Topisirovic and Sonenberg, 2011). These proteins are active when phosphorylated; therefore, a larger ratio of phosphorylated protein/total protein indicates increased activity of these regulatory proteins and more downstream protein synthesis. Activity-regulated cytoskeleton-associated protein (ARC) expression was measured as a downstream indicator of increased translation. Arc is an immediate early gene; its expression is correlated with neural activity (Lyford et al., 1995).

The effects of the drugs on activating translation were evaluated in both rat and mouse acute hippocampal slices. Treatment chambers for the slices were constructed from 40 mL specimen containers (Simport Scientific), Netwell 24 mm mesh inserts (Electron Microscopy Sciences), and small, sterilizer glass beads (Sigma). Tubing with flow regulators from standard, disposable intravenous sets delivered 95% O_2_/ 5% CO_2_ to the bottom of each chamber.

Two-month-old rats were decapitated to obtain acute hippocampal slices (400 µm). Brains were extracted and immersed in chilled, oxygenated cutting medium (220 mM sucrose, 20 mM NaCl, 2.5 mM KCl, 1.25 mM NaH2PO4, 26 mM NaHCO3, 10 mM glucose, 2 mM ascorbic acid, and 2.5 mM MgSO4). Hippocampi were bilaterally dissected in cold cutting medium. Using a dry McIlwainTM Tissue Chopper (Stoelting Co), 400 µm hippocampal slices were cut and stabilized in chambers containing cutting medium bubbled with 95% O_2_, 5% CO_2_ for 20 minutes at 35 °C. Slices were transferred and pre-incubated in oxygenated (95% O_2_, 5% CO_2_) artificial cerebrospinal fluid (aCSF; 125 mM NaCl, 3 mM KCl, 1.25 mM NaH_2_PO_4_, 26 mM NaHCO_3_, 2 mM ascorbic acid, 10 mM glucose, 1.5 mM MgSO_4_, and 2.5 mM CaCl_2_) at 34°C for 1 h.

Wild-type, male C57/BL6J mice, 12-13 weeks-old, were decapitated to obtain acute 400 µm hippocampal slices prepared similar to those from rats. Slices were transferred to oxygenated aCSF (124 mM NaCl, 2.5 mM KCl, 1.25 mM NaH2PO4, 24 mM NaHCO3, 2 mM ascorbic acid, 10 mM glucose, 1.5 mM MgSO4, and 2.5 mM CaCl_2_) at 35°C before adding pharmacological agents (Zhou and Baudry, 2006).

Drugs were added directly into aCSF for 1 h at the indicated final concentrations: 10 μM PCP, 100 µM (S)-3,5-DHPG, 25 µM MK801 maleate, 50 µM AP5, 0.5 µM Ro 25-6981 maleate, and 0.5 μM NVP-AAM077. Slices co-treated with DHPG received pre-treatment for 10 minutes with DHPG followed by addition of the NMDAR antagonist for 1 h.

At the end of the drug and control treatments, slices were collected in 4°C lysis buffer containing protease and phosphatase inhibitors (65 mM Tris-HCl, 10% glycerol, 2% SDS, 5 mM EGTA, 5 mM EDTA, 200 µM Na_3_VO_4_, 200 µM PMSF, 2 mM NaF, 0.5% Triton, 10 mM Beta-glycerophosphate disodium pentahydrate, 10 mM sodium pyrophosphate, phosphatase inhibitor cocktail (Roche, Cat.No.4906837001) and protease inhibitor cocktail (Roche, Cat.No.4693124001) as described previously (Jourdi et al., 2009). Tissue was mechanically homogenized, dry sonicated (Branson Sonifier 250), and then centrifuged at 10,000g x 2 min. The supernatant was denatured for 2 rounds x 5 min in a 95°C heat block. Protein determination was performed with the Micro BCA^TM^ Protein Assay Kit (Pierce, Cat. No.23235). Sample concentrations were adjusted with lysis buffer and then denatured at 95°C for 5 min in 20% loading buffer (10% SDS, 0.3125M Tris HCl pH 6.8, 3.575M beta-mercaptoethanol, 0.05% bromophenol blue in H_2_O). Forty µg denatured protein per well was resolved by 8-12% SDS-PAGE homemade gels in parallel with multi-colored molecular weight marker (Bio-Rad Laboratories, Cat.No.1610395). Gels were run at 100V for 1-2 h and transferred overnight at 8- 12V, 4°C onto 0.2uM PVDF membranes (GE Healthcare Biosciences). After transfer, gels were stained for 1 h with Coomassie Blue R-250 0.5% and membranes were stained with MemCode^TM^ Reversible Stain (Pierce, Cat.No.24585) to examine the quality of the protein transfer. Membranes were blocked for 1 h in I-Block^TM^ (Applied Biosystems) protein-based blocking reagent and then incubated with primary antibodies overnight at 4°C. Between primary and secondary antibody incubations, membranes were rinsed three times in 1X Tris buffer solution (TBS) and then washed in I-Block^TM^ 3 x 5 min. Membranes were incubated for 1 h at room temperature in horseradish peroxidase-conjugated secondary antibodies in I-Block^TM^ solution, followed by three rinses and 3 x 5 min washes in 1X TBS. Imaging and detection were performed with ECL Select^TM^ (GE Healthcare, Cat.No.RPN2235) on a C-DiGit^TM^ Blot Scanner (LI-COR, Cat.No.3600-00). Images were analyzed and presented as before (Jourdi et al., 2009).

#### Antibodies

Primary antibodies were phospho-mTOR (Santa Cruz Biotechnology, Cat.No.101738, 1:1000), mTOR (Pierce, Cat.No.PA534663, 1:1000), ARC (Cedarlane-Synaptic Systems, Cat.No.156003(SY) 1:1000), phospho-AKT (Cell Signaling Technology, Cat.No.9271S, 1:1000), AKT (Cell Signaling Technology, Cat.No.9272S, 1:1000), phospho-ERK1/2 (Cell Signaling Technology, Cat.No.9101S, 1:1000), ERK1/2 (Cell Signaling Technology, Cat.No.9102S, 1:1000), phospho-4EBP1 (Cell Signaling Technology, Cat.No.9459S, 1:1000), 4EBP1 (Cell Signaling Technology, Cat.No.9452S, 1:1000). Actin (Sigma, Cat.No.A2228, 1:20000) and GAPDH (Abcam, Cat.No.ab8245, 1:10000) were used as loading controls. Secondary antibodies were goat anti-rabbit (Pierce, Cat.No.31462, 1:10000) and goat anti-mouse (AnaSpec, Cat.No.28173, 1:10000). All antibodies were diluted in I-Block^TM^ protein-based blocking reagent (Applied Biosystems, Cat.No.T2015).

Blots were reacted with primary antibodies against the following proteins to monitor activation of the translation machinery: 4-EBP1, phospho-4-EBP1, protein kinase B (AKT), phospho-AKT, the mechanistic target of rapamycin (mTOR), phospho-mTOR, extracellular-signal-regulated kinase (ERK)1/2 and phospho-ERK1/2 and against the activity-regulated cytoskeleton-associated protein (ARC) to monitor protein synthesis. All blots were reacted with GAPDH and actin as loading controls. Results from all western blots with phospho-specific antibodies were normalized against the expression levels of the corresponding native proteins, whose expression levels were not changed by the PCP or DHPG treatments. ARC protein levels were normalized against GAPDH or actin, which were used as loading controls. Data were calculated as percent change compared to the control condition from six independent experiments.

### Statistical Analyses

Comparisons were performed by Student’s t test, one-way, and two-way ANOVAs, as appropriate. When needed, Tukey’s and Dunnett’s post-hoc comparisons were performed. Statistical significance was accepted when p < 0.05.

## AUTHOR CONTRIBUTIONS

Data were collected by EHP (awake electrophysiology), BR (electrophysiology under anesthesia), MvD (*ex vivo* electrophysiology), EW and MT-R (mouse behavior), KWT, JG, and HJ (biochemistry); data were analyzed by H-YK, MvD, HJ, EL, SCS, and AAF; HJ, JMA and AAF supervised research; AAF designed research; AAF wrote the paper and all authors contributed to editing and figures.

## ACKNOWLEDGEMENTS

Supported by NIMH grants R21MH082417, R01MH084038, and R01NS105472.

